# Dietary choline intake is necessary to prevent systems-wide organ pathology and reduce Alzheimer’s disease hallmarks

**DOI:** 10.1101/2022.08.14.503929

**Authors:** Nikhil Dave, Jessica M. Judd, Annika Decker, Wendy Winslow, Patrick Sarette, Oscar V. Espinosa, Jessica Sandler, Alina Bilal, Savannah Tallino, Ian McDonough, Joanna K Winstone, Erik A. Blackwood, Christopher Glembotski, Timothy Karr, Ramon Velazquez

**Author notes:** **Correspondence:** Ramon Velazquez Ph.D., Neurodegenerative Disease Research Center, Arizona State University, 797 E Tyler St, Tempe, AZ 85287. First co-authors (Nikhil Dave, Jessica M. Judd).

## Abstract

Evidence suggests that environmental factors may contribute to Alzheimer’s disease (AD). The B-like vitamin choline plays key roles in body- and brain-related functions. Choline produced endogenously by the phosphatidylethanolamine N-methyltransferase (PEMT) enzyme in the liver is not sufficient for adequate physiological functions, necessitating daily dietary intake. ∼90% of Americans don’t reach the recommended daily choline intake. Thus, it’s imperative to determine whether dietary deficiency increases disease outcomes. Here, we placed 3xTg-AD, a model of AD, and non-transgenic (NonTg) control mice on either a sufficient choline (ChN) or choline deficient (Ch-; choline deficiency) diet from 3 to 12 (early to late adulthood) months of age. Ch- reduced plasma choline and acetylcholine levels, increased weight, and impaired both glucose metabolism and motor function in NonTg, with 3xTg-AD mice showing greater deficits. Tissue analyses showed cardiac and liver pathology, and elevated Amyloid-β and phosphorylated tau in the hippocampus and cortex of 3xTg-AD Ch- mice. Unbiased proteomic analyses revealed Ch- altered hippocampal networks associated with microtubule function and postsynaptic membrane regulation. In plasma, Ch- altered protein networks associated with insulin metabolism, mitochondrial function, and inflammation. Collectively, our data highlight that dietary choline intake is necessary to prevent systems-wide organ pathology and reduce AD hallmark pathologies.

## Introduction

Alzheimer’s disease (AD) is one of the most prevalent age-related neurodegenerative disorders in the US, with more than 6 million Americans currently living with the disease and a projected 16 million affected by 2050 (1). AD is characterized by two hallmark neuropathologies: extracellular amyloid-beta (Aβ) plaques and intracellular neurofibrillary tangles (NFT), resulting in a progressive loss of cognitive abilities and memory (2). Other pathologies also occur in AD cases, including neuroinflammation (3, 4), cardiac pathology (5), insulin dysregulation (4), and energy dyshomeostasis (4, 6). These physiological alterations show that AD is not limited to neuropathology – rather, it is a complex, systems-wide disease affecting several different metabolic and cellular processes throughout the human body. While a wealth of literature investigates these disease parameters separately, it is still unclear how exactly they contribute to AD pathogenesis or when they chronologically coincide with Aβ and NFT pathology. Notably, while genetic mutations contribute to familial AD, which accounts for < 5% of those affected, an abundance of work has highlighted that environmental factors, including diet, may play a role in sporadic AD, which accounts for > 95% of total AD cases (4, 5, 7, 8).

Choline, an essential nutrient found in a variety of foods, is a critical part of the metabolic pathway responsible for the creation of choline phospholipids, betaine, and acetylcholine (ACh), a key neurotransmitter involved in neurogenesis, synapse formation, learning, and memory (9). Betaine serves as a methyl group donor in the conversion of homocysteine to methionine (9), an amino acid that contributes to epigenetic mechanisms regulating gene expression (10). While choline is produced endogenously by the phosphatidylethanolamine N-methyltransferase (PEMT) protein in the livers of both mice and humans, endogenous production is not sufficient for normal metabolic functioning, requiring dietary intake (11). In 1998, the Institute of Medicine (IOM) established an adequate daily intake (ADI) threshold for choline consumption to prevent fatty liver disease, stating that adult men and women should consume 550 mg/day and 425mg/day, respectively (12). Pregnant women should consume 550 mg/day for healthy fetal development (9). However, recent studies have shown that ∼90% of the US population is deficient in dietary choline (13).

Significant evidence shows that choline is important for healthy brain function. Maternal choline supplementation (MCS) of 4.5 times the ADI has been shown to produce important cognitive benefits for offspring: these findings have been corroborated in mouse studies of AD and Down syndrome (7–9, 14). More recently, adulthood choline supplementation in a mouse model of AD has been shown to significantly reduce Aβ plaque density, learning and memory deficits, and brain inflammation (8). Additional work has highlighted the relationship between choline and dysfunction of systems-wide cellular and molecular processes that are also implicated in AD. For example, choline supplementation is associated with attenuated microglial activation and reduced insulin resistance in the brain and periphery (8, 15), and choline deficiency plays a role in cardiovascular disease, liver toxicity, and hypertension (12, 16); perturbations to these same peripheral systems are known risk factors for AD, suggesting that choline deficiency could be a shared mechanism of all these ailments. Moreover, previous work has shown that a functional single nucleotide polymorphism (rs7964) in the gene encoding PEMT is associated with AD (17), suggesting that abnormalities in endogenous choline production may be associated with increased AD incidence. A recent report showed that choline restores lipid homeostasis deficits in human IPSC-derived astrocytes from apolipoprotein E (*APOE)* 4 positive patients; *APOE4* is also a significant risk factor for AD (18). Taken together, these studies suggest that reduced levels of choline may elevate the risk of AD, highlighting the importance of adequate intake.

Despite a myriad of publications demonstrating the positive effects of choline supplementation, there is a limited understanding of the health and cognitive impacts of dietary choline deficiency throughout adulthood. Here, we sought to elucidate the effects of dietary choline deficiency in healthy aging and in AD, placing NonTg mice and a mouse model of AD (3xTg-AD) on a choline-deficient diet throughout adulthood. We hypothesized that a choline deficient diet in adulthood will induce system-wide cellular and molecular dysfunction throughout the body and brain, increasing the risk of AD across several pathogenic axes.

## Results

At three months of age, we exposed female NonTg and 3xTg-AD mice to either a control choline (ChN; 2.0g/kg; #TD.180228) or choline deficient (Ch-; choline deficiency; 0.0g/kg; #TD.110617; Supplementary Fig. 1A, B) diet. All other diet components were identical. All mice were aged to 10 months and tested in the rotarod task to assess motor function, and the Morris water maze (MWM) to assess spatial reference learning and memory (Fig. 1A). There were four experimental groups (NonTg ChN, n = 20; 3xTg-AD ChN, n = 15; NonTg Ch-, n = 18; 3xTg-AD Ch-, n = 16; Fig. 1B). After testing, all mice were sacrificed and their blood, hearts, livers, and brains were collected for pathological and proteomic assessment.

**Figure 1.**
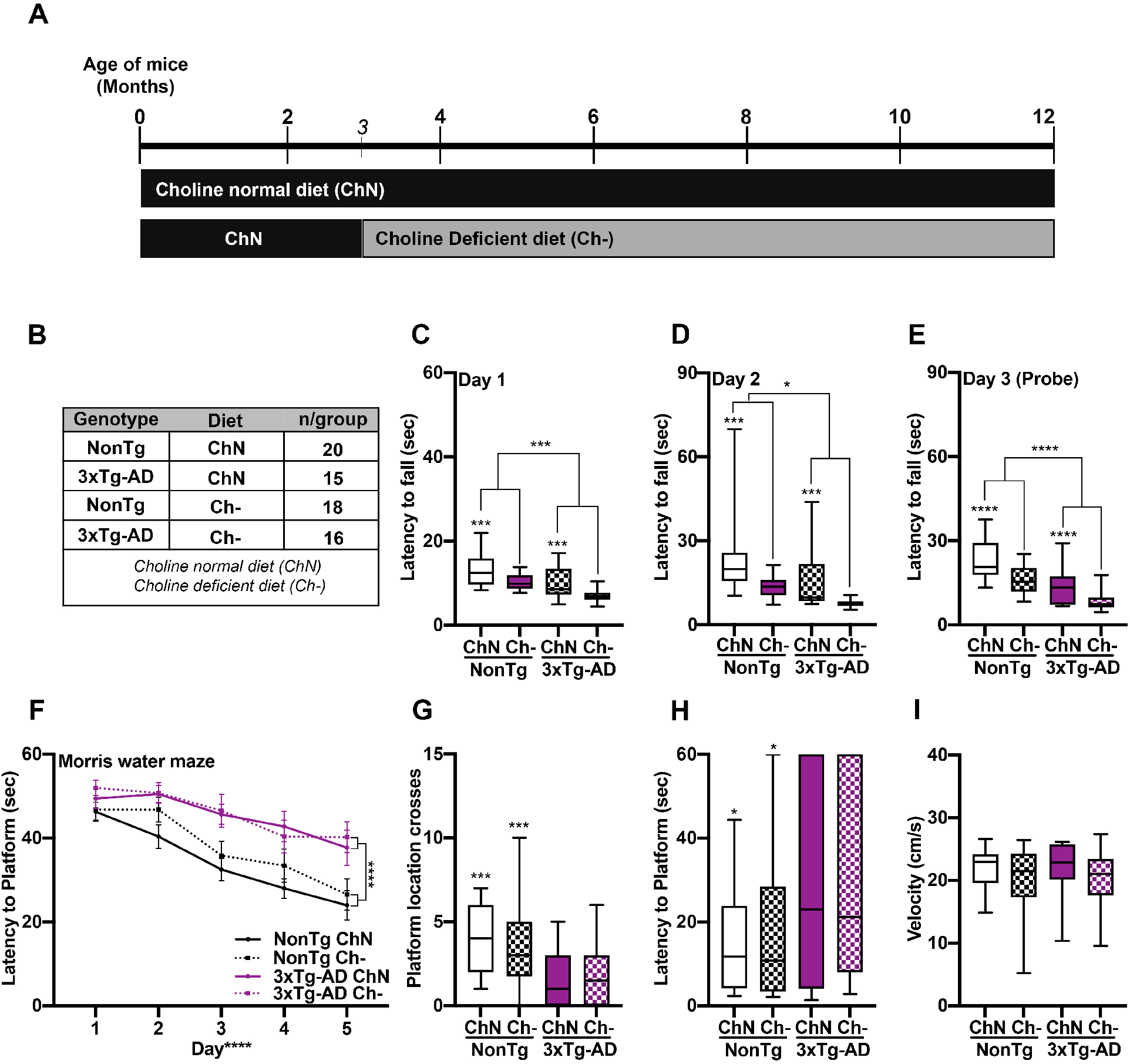
Ch- impairs motor function in both NonTg and 3xTg-AD mice. **A**. Experimental design. **B**. Groups and sample size. **C-E**. Motor function assessment via the rotarod task illustrated significant main effects of genotype, where the 3xTg-AD mice fell of the spinning rod significantly faster than NonTg mice on Day 1 (p = 0.001), Day 2 (p = 0.016) and the Day 3 probe trial (p < 0.0001). Assessment also showed significant main effects of diet, where the Ch- mice fell significantly faster on Day 1 (p = 0.0002), Day 2 (p = 0.0002) and Day 3 (p < 0.0001), than their ChN counterparts. **F**. Morris water maze testing revealed significant effects of genotype, where the 3xTg-AD had a higher latency to find the platform across the 5-days of learning than the NonTg mice (p < 0.0001). **G-I**. During the Day 6 probe trial, the 3xTg-AD mice crossed the platform significantly fewer times (p = 0.0008) and had a higher latency to first cross the platform location (p = 0.0105) than the NonTg mice. Swim speed was equal across the groups. For box plots, the center line represents the median value, the limits represent the 25^th^ and 75^th^ percentile, and the whiskers represent the minimum and maximum value of the distribution. Line graphs are mean ± SE. * p < 0.05, *** p < 0.001, **** p < 0.0001.

### Ch- throughout adulthood impairs motor function

Motor function was assessed using the rotarod task. During the first two training days, we found significant main effects of genotype and diet for latency to fall off the spinning rod on Day 1 (F(1, 65) = 16.72, p = 0.001; Fig. 1C), (F_(1, 65)_ = 15.86, p = 0.0002) and Day 2 (F_(1,65)_ = 6.114, p = 0.016; Fig. 1D), (F_(1, 65)_ = 15.69, p = 0.0002), where 3xTg-AD mice fell sooner than the NonTg mice, and Ch- mice fell sooner than the ChN mice. On Day 3, we performed six probe trials and found highly significant main effects of genotype (F_(1, 65)_ = 20.99, p < 0.0001; Fig. 1E) and diet (F_(1, 65)_ = 20.56, p < 0.0001), where 3xTg-AD mice fell sooner than the NonTg mice, and Ch- mice fell sooner than their ChN counterparts. These results indicate that the AD phenotype and adulthood Ch- is sufficient to impair motor function.

Mice were then tested in the MWM for 6 consecutive days. One 3xTg-AD Ch- mouse was excluded due to an inability to swim. During the first 5 training days, mice received four trials/day. For latency to find the hidden platform, we found a significant main effect of day, indicating learning across all groups (F_(1, 256)_ = 26.932 p < 0.0001; Fig. 1F). We also found a significant main effect of genotype (F_(1, 256)_ = 23.335, p < 0.0001), where the 3xTg-AD mice took significantly longer to find the hidden platform compared to the NonTg mice. No significant main effects or interactions were found for diet. On Day 6, the platform was removed, and mice were tested in a 60-s probe trial to assess spatial memory. 3xTg-AD mice crossed the platform location significantly fewer times than the NonTg mice (F_(1, 64)_ = 12.276, p = 0.0008; Fig. 1G) and had a higher latency to first cross the platform location (F_(1, 64)_ = 6.948, p = 0.0105; Fig. 1H), illustrating spatial memory deficits. No diet effects were detected. Swim speed during the probe trials was similar across groups (Fig. 1I). These data indicate that Ch- does not exacerbate spatial memory impairments in the 3xTg-AD mice nor impair performance in NonTg mice.

### Ch- increases weight, induces glucose metabolism impairments, and reduces plasma choline and ACh levels in NonTg and 3xTg-AD mice

Choline plays an essential role in glucose metabolism, another biological system that is dysfunctional in AD (19). Thus, we examined body weight and performed a glucose tolerance test (GTT). For percent weight change from baseline, we found significant main effects of genotype (F_(1, 65)_ = 51.96, p < 0.0001) and diet (F_(1, 65)_ = 52.86, p < 0.0001), where 3xTg-AD mice had a higher percent weight change than NonTg mice. Ch- mice had a higher percent weight change than their ChN counterparts (Fig. 2A, B). We also found a significant genotype by diet interaction (F_(1, 65)_ = 10.91, p = 0.0016). Post hoc analysis revealed that the NonTg Ch- mice had a significantly higher percent weight change from baseline than their ChN counterparts (p = 0.026). 3xTg-AD Ch- mice showed a higher percentage weight change from baseline than 3xTg-AD ChN mice (p < 0.0001). Notably, NonTg Ch- and 3xTg-AD ChN mice showed no significant difference (p > 0.99), indicating the weight gain in NonTg Ch- mice phenocopies that seen in the AD mouse. To determine if body weight differences arose from increased food intake, we performed a six-day food consumption test. We found a significant main effect of genotype (F_(1, 14)_ = 11.69, p = 0.0042; Fig. 2C), where the 3xTg-AD mice consumed significantly more than NonTg mice, but no significant diet effects, illustrating that body weight differences between diets (Fig. 2D) could not be attributed to food consumption.

**Figure 2.**
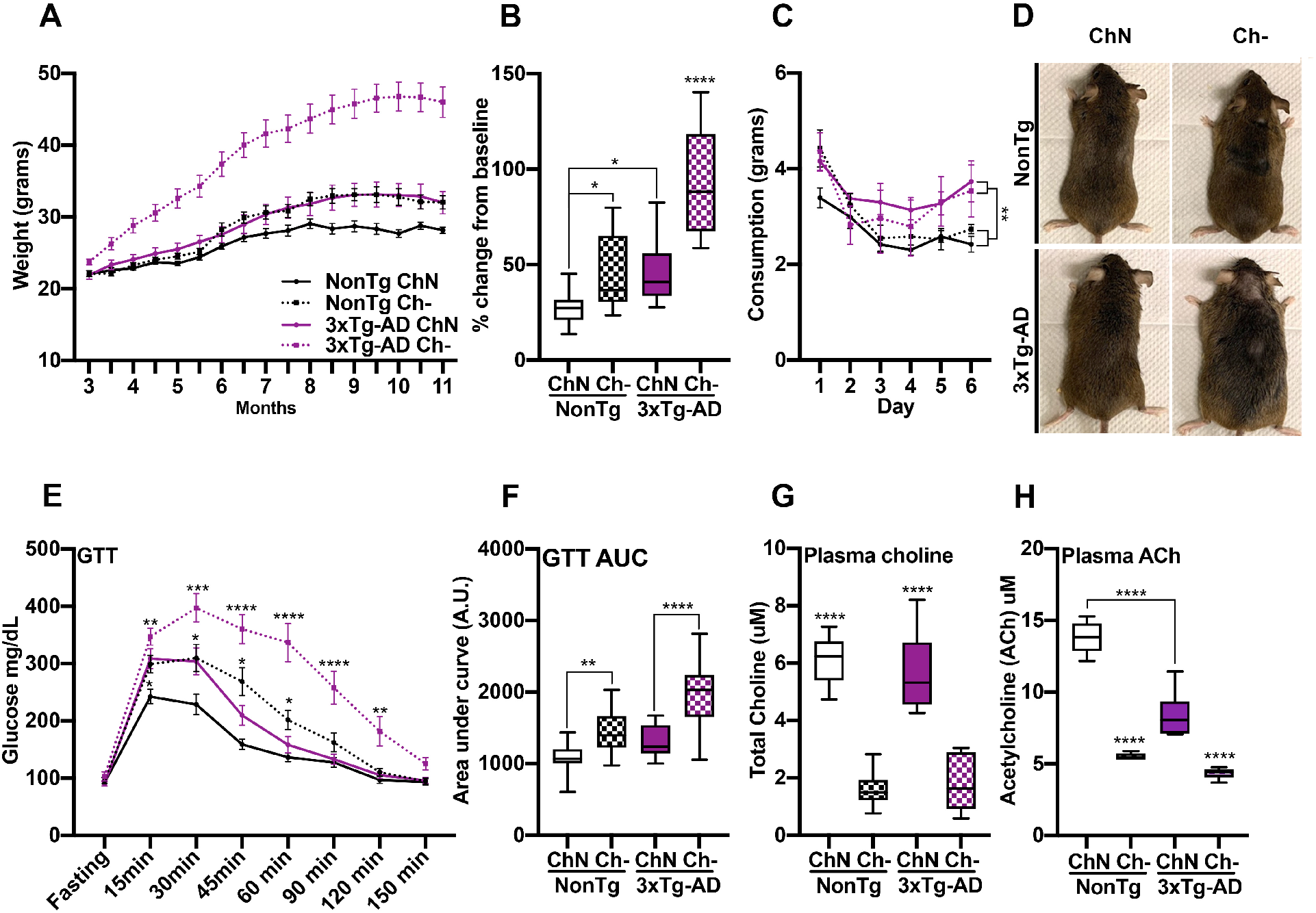
Ch- promotes weight gain, impairs glucose metabolism, and reduces plasma choline and acetylcholine levels. **A**. Body weight across age. **B**. Percent weight change from baseline shows elevated levels in the NonTg Ch- mice compared to NonTg ChN mice (p= 0.026) and in the 3xTg-AD Ch- mice compared to 3xTg-AD ChN mice (p < 0.0001). **C**. 3xTg-AD mice had significantly higher food intake across the six days of the food consumption test than the NonTg mice (p = 0.0042), but no diet effect was found. **D**. Representative images of mice illustrating weight differences. **E**. 3xTg-AD Ch- mice had significantly higher glucose levels in the glucose tolerance test (GTT) from the 15 through 150 min timepoints than their ChN counterparts (p < 0.05). NonTg Ch- mice had higher glucose levels than their ChN counterparts at the 15 through 60 min time points (p < 0.05). **F**. GTT Area under the curve (AUC) analysis showed higher levels in NonTg Ch- mice than their ChN counterparts (p = 0.0042), and 3xTg-AD Ch- mice had higher levels than their ChN counterparts (p < 0.0001). **G**. Ch- significantly reduced plasma choline levels (p < 0.0001). **H**. Ch- significantly reduced acetylcholine (ACh) levels (p < 0.0001). 3xTg-AD ChN mice had significantly lower ACh levels than the NonTg ChN mice (p < 0.0001). Line graphs are mean ± SE. For box plots, the center line represents the median value, the limits represent the 25^th^ and 75^th^ percentile, and the whiskers represent the minimum and maximum value of the distribution. * p < 0.05, ** p < 0.01, *** p < 0.001, **** p < 0.0001.

In the GTT, we found significant main effects of genotype (F_(1, 65)_ = 26.02, p < 0.0001) and diet (F_(1, 65)_ = 47.01, p < 0.0001), where the 3xTg-AD mice show higher glucose levels than NonTg mice. Ch- mice had higher glucose levels than the ChN mice (Fig. 2E). We also found a significant genotype by diet by time interaction (F_(7,455)_ = 2.674, p = 0.0101). Post hoc analysis revealed that 3xTg-AD Ch- mice had significantly higher glucose levels from the 15 through 150-minute time points compared to their ChN counterparts (p < 0.05). Strikingly, NonTg Ch- mice had higher glucose levels than their ChN counterparts at the 15 through 60 min time points (p < 0.05). Lastly, we analyzed glucose area under the curve (AUC), as it provides a better assessment of glucose tolerance (6). We found significant main effects of genotype (F_(1, 65)_ = 26.04, p < 0.0001) and diet (F_(1, 65)_ = 47.25, p < 0.0001), where the glucose AUC for 3xTg-AD mice was higher than for the NonTg mice and higher for the Ch- mice than for the ChN mice (Fig. 2F). We also found a significant genotype by diet interaction (F_(1, 65)_ = 4.393, p = 0.040). Post hoc analysis revealed that AUC was higher in NonTg Ch- than their ChN counterparts (p = 0.0042), and in 3xTg-AD Ch- compared to the 3xTg-AD ChN mice (p < 0.0001). AUC was similar between the NonTg Ch- and 3xTg-AD ChN mice (p > 0.9999). Collectively, these results show that Ch- increases weight and impairs glucose metabolism in both the control and AD mice, with the 3xTg-AD mice showing greater deficits. This is notable because weight gain and impaired glucose metabolism are risk factors for AD (6), highlighting the importance of dietary choline to deter metabolic deficits.

To determine whether choline levels were affected by diet, we collected blood at euthanasia and isolated plasma. We found a significant main effect of diet (F_(1, 20)_ = 96.08, p < 0.0001), where the Ch- mice (n = 6/group) had significantly reduced levels of circulating choline in their plasma than the ChN mice had (Fig. 2G). Next, we measured plasma ACh levels and found a significant main effect of diet (F_(1, 20)_ = 222.80, p < 0.0001), where the Ch- mice had lower levels than their ChN counterparts and a significant genotype by diet interaction (F_(1,20)_ = 26.00, p < 0.0001; Fig. 2H). Post hoc analysis revealed that the 3xTg-AD ChN mice had significantly reduced levels of plasma ACh than the NonTg ChN mice.

### Ch- induces cardiac and liver pathology in NonTg and 3xTg-AD mice

Given the role of choline in cardiac dysfunction and previous reports of cardiac dysfunction in AD (5, 16), we examined cardiac pathology. We first measured heart weight normalized by tibia length as a measure of pathological cardiac hypertrophy (20); NonTg ChN (n = 9), NonTg Ch- (n = 7), 3xTg-AD ChN (n = 4), and 3xTg-AD Ch- (n = 6). We found significant main effects of genotype (F_(1,22)_ = 29.63, p < 0.0001) and diet (F_(1,22)_ = 13.78, p = 0.001), where 3xTg-AD mice had a higher heart weight than NonTg mice. Ch- groups had higher heart weights than the ChN mice (Fig. 3A). We also found a significant genotype by diet interaction (F_(1,22)_ = 13.78, p = 0.001). Post hoc analysis revealed that the 3xTg-AD Ch- mice had a higher heart weight than the other groups (p = 0.0018). Next, total RNA was extracted from snap-frozen left ventricular extracts and subjected to qRT-PCR for transcript analysis; NonTg ChN (n = 6), NonTg Ch- (n = 6), 3xTg-AD ChN (n = 4), and 3xTg-AD Ch- (n = 6). We examined the expression levels of *Col1a1, Myh7*, and *Nppa*, which are all genes whose increased expression is associated with cardiac pathology (21–23). We found significant genotype by diet interactions ((F_(1,18)_ = 27.85, p < 0.0001; Fig. 3B), (F_(1,18)_ = 9.743, p = 0.006), (F_(1,18)_ = 19.17, p = 0.0004), respectively). Post hoc analysis revealed that the NonTg ChN mice had lower expression of *Colla1, Myh7*, and *Nppa* than all other groups (p < 0.05). These data demonstrate that cardiac dysfunction can be caused by both AD pathogenesis and Ch-.

**Figure 3.**
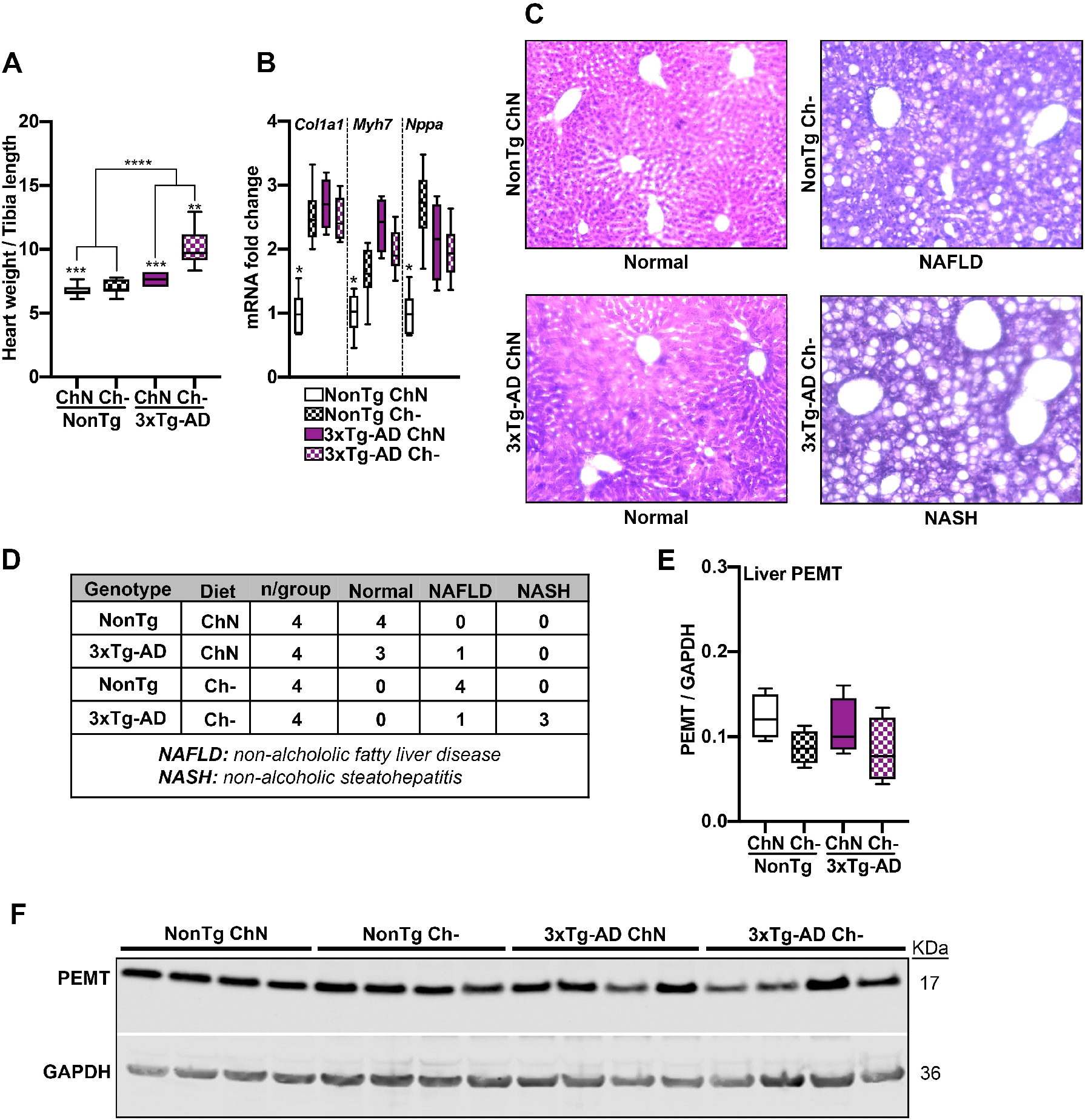
Ch- induced liver and cardiac pathology in NonTg and 3xTg-AD mice. **A**. 3xTg-AD mice had a higher heart weight than NonTg mice (p < 0.0001), and the Ch- groups had higher heart weights than the ChN mice (p = 0.001). 3xTg-AD Ch- mice had the highest heart weight (p = 0.0018). **B**. mRNA expression levels of *Col1a1, Myh7*, and *Nppa* were elevated in 3xTg-AD ChN mice and both NonTg and 3xTg-AD Ch- mice compared to the NonTg ChN group (p < 0.05). **C**. Photomicrographs of liver tissue stained with Hematoxylin and Eosin (H&E). **D**. Table reporting liver pathology in Ch- mice. **E, F**. Western blot and quantification for PEMT illustrates no difference across groups. For box plots, the center line represents the median value the limits represent the 25^th^ and 75^th^ percentile, and the whiskers represent the minimum and maximum value of the distribution. * p < 0.05, ** p < 0.01, *** p < 0.001, **** p < 0.0001.

The existing guidelines for dietary choline established in 1998 by the IOM were developed to prevent non-alcoholic fatty liver disease (NAFLD; 12). To determine if adulthood Ch- throughout life induced liver pathology, we acquired liver tissue from mice at death (n = 4 mice/group) and assessed pathology using a Hematoxylin and Eosin (H&E) stain (Fig. 3C). Staining for the ChN mice showed that NonTg mice had healthy livers and one 3xTg-AD mouse showed evidence of NAFLD (Fig. 3D). All the NonTg Ch- mice exhibited NAFLD, and three 3xTg-AD mice showed non-alcoholic steatohepatitis (NASH). We measured protein levels of PEMT from liver tissue to determine if Ch- led to compensatory upregulation. We found no significant differences across the four graphs (Fig. 3E, F). Collectively, these results show that a Ch- diet induces pathology in multiple organ systems associated with metabolic function and that this dysfunction may potentially contribute to AD-pathogenesis.

### Ch- reduces hippocampal and cortical choline levels and exacerbates soluble and insoluble Amyloid-ß (Aß) pathology and tau hyperphosphorylation

We measured hippocampal and cortical (n = 6/group) levels of choline to determine the effects of Ch-. We found a significant main effect of diet for both hippocampal (F_(1, 20)_ = 133.50, p < 0.0001) and cortical (F_(1, 20)_ = 165.30, p < 0.0001) choline levels, where the Ch- mice had significantly lower levels than the ChN mice (Fig. 4A, B). To understand the effects of Ch- on AD pathogenesis, we used ELISAs to quantify soluble and insoluble Aß in 3xTg-AD Ch- (n = 8) and ChN (n = 7) mice. NonTg mice do not display Aß or tau pathology and therefore were excluded from these analyses (8). The two major isoforms of Aß are Aß_40_ and Aß_42_, found in both soluble and insoluble fractions (2). These isoforms of Aß aggregate into Aß oligomers, which are neurotoxic precursors for insoluble Aß plaques (24). For soluble Aß_40_ fractions, we found no significant differences in hippocampal (t_(13)_ = 1.712, p = 0.111) or cortical (t_(13)_ = 1.685, p = 0.116) levels between 3xTg-AD ChN and Ch- mice (Fig. 4C). For soluble Aß_42,_ we found a significant difference where 3xTg-AD Ch- mice had higher levels in the hippocampus (t_(13)_ = 2.833, p = 0.014) and cortex (t_(13)_ =11.59, p < 0.0001) compared to the ChN counterparts (Fig. 4D). Similarly, we found a significant difference in toxic soluble Aß oligomers, where 3xTg-AD Ch- mice had higher levels in the hippocampus (t_(13)_ = 3.335, p = 0.0054) and cortex (t_(13)_ = 2.648, p = 0.021) compared to their ChN counterparts (Fig. 4E). For insoluble Aß_40_ fractions in the hippocampus, we found a non-significant trend where 3xTg-AD Ch- had higher levels than the ChN mice (t_(13)_ = 1.919, p = 0.077; Fig. 3F). In the cortex (t_(13)_ = 2.398, p = 0.032), we found that 3xTg-AD Ch- mice exhibited higher levels of insoluble Aß_40_ compared to their ChN counterparts (Fig. 4F). For insoluble Aß_42_ fractions, we found a significant difference where 3xTg-AD Ch- mice had higher levels in the hippocampus (t_(13)_ = 2.357, p = 0.035) and cortex (t_(13)_ = 13.90, p < 0.0001) compared to their ChN counterparts (Fig. 4G). Lastly, we stained tissue with Thioflavin S to detect Aß sheets and found increased expression in the entorhinal cortex of 3xTg-AD Ch- mice (Fig. 4H). These results illustrate that Ch- throughout adulthood increases the levels of toxic Aß pathology in 3xTg-AD mice.

**Figure 4.**
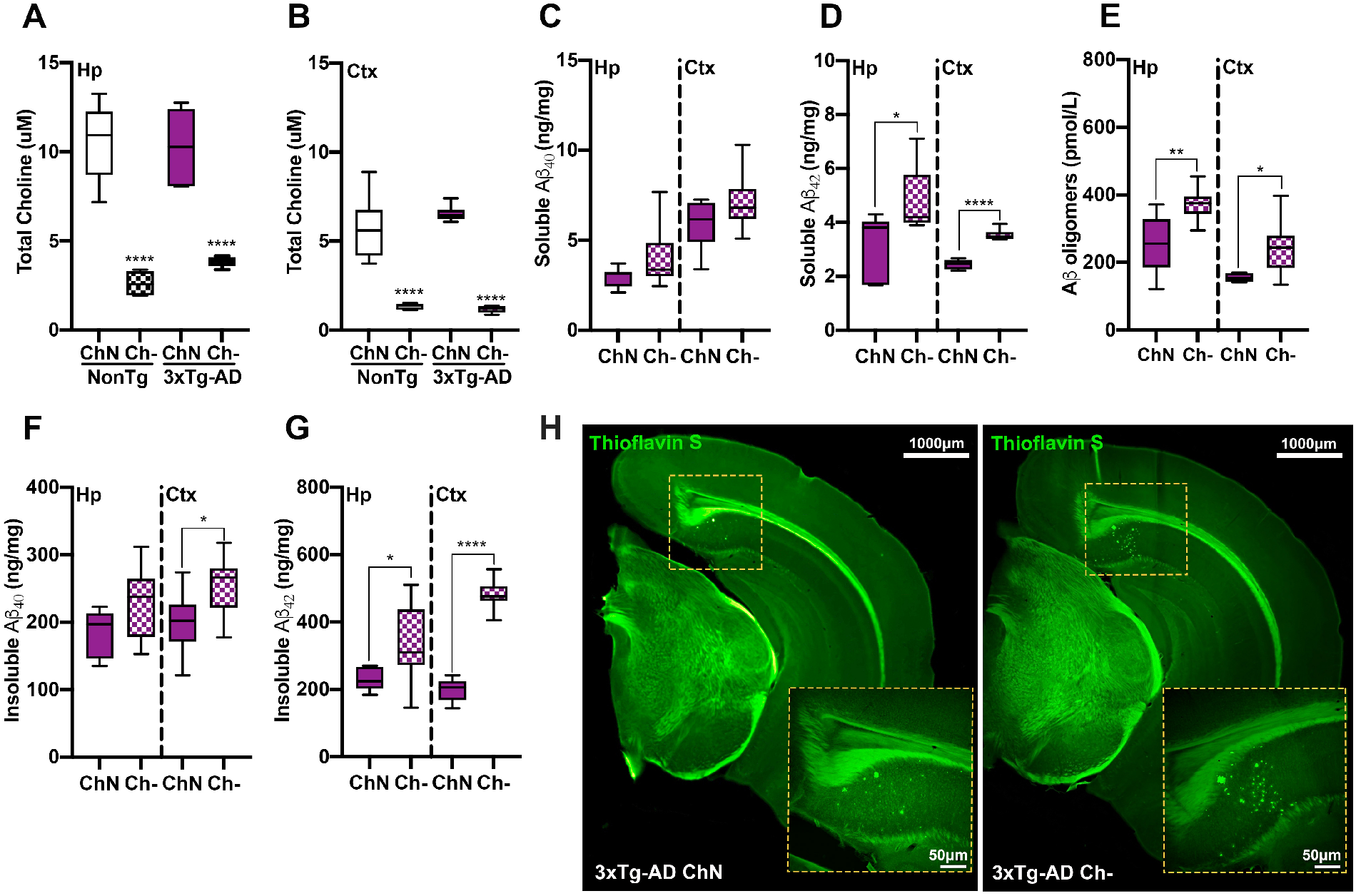
Ch- reduces hippocampal and cortical choline levels and increases soluble and insoluble Amyloid-ß fractions. **A, B**. Hippocampal (Hp; p < 0.0001) and cortical (Ctx; p < 0.0001) choline levels were significantly reduced in the Ch- mice compared to their ChN counterparts. NonTg mice do not display Aß pathology and were therefore excluded from Aß analyses. **C**. No significant differences were detected in soluble Aß_40_. **D**. Soluble Aß_42_ levels were significantly elevated in the Hp (p = 0.014) and Ctx (p < 0.0001) of 3xTg-AD Ch- mice. **E**. Aß oligomer levels were significantly elevated in the Hp (p =0.0054) and Ctx (p = 0.021) of 3xTg-AD Ch- mice. **F**. For insoluble Aß_40_ levels, we found a non-significant trend in the Hp (p = 0.077) and significantly elevated levels in the Ctx (p = 0.032) of 3xTg-AD Ch- mice. **G**. Insoluble Aß_42_ levels were significantly elevated in the Hp (p = 0.035) and Ctx (p < 0.0001) of 3xTg-AD Ch- mice. **H**. Photomicrographs of Thioflavin S staining illustrating Aß-plaques in the entorhinal cortex. For box plots, the center line represents the median value, the limits represent the 25^th^ and 75^th^ percentile, and the whiskers represent the minimum and maximum value of the distribution. * p < 0.05, ** p < 0.01, *** p < 0.001, **** p < 0.0001.

We also sought to understand the effects of Ch- on tau pathogenesis. We used ELISAs to detect hippocampal and cortical soluble and insoluble fractions of phosphorylated (p) Tau at serine 181 (pTau Ser181) and serine 396 (pTau Ser396) - two sites known to be associated with tau pathogenesis in AD (25, 26). For soluble pTau Ser181, we found a significant difference where 3xTg-AD Ch- mice had higher levels in the hippocampus (t_(13)_ = 9.045, p < 0.0001) and cortex (t_(13)_ = 7.691, p = 0.0009) compared to their ChN counterparts (Fig. 5A). For soluble pTau Ser396, we found a significant difference where 3xTg-AD Ch- mice had higher levels in the hippocampus (t_(13)_ = 13.92, p < 0.0001) and cortex (t_(13)_ = 4.245, p < 0.001) compared to their ChN counterparts (Fig. 5B). For insoluble pTau Ser181, we found a significant difference where the 3xTg-AD Ch- mice had higher levels in the hippocampus (t_(13)_ = 2.228, p = 0.0442) and cortex (t_(13)_ = 3.356, p = 0.0052) than their ChN counterparts (Fig. 5C). No significant differences between 3xTg-AD ChN and Ch- mice were detected for insoluble pTau Ser396 in hippocampal (t_(13)_ = 0.622, p = 0.545; Fig. 3I) or cortical (t_(13)_ = 1.436, p = 0.175) tissue (Fig. 5D). Lastly, we stained against AT8 (Ser202/Thr205), which is associated with intraneuronal tau filaments (27). AT8-positive cells were significantly higher in 3xTg-AD Ch- mice (t_(6)_ = 2.462, p = 0.049; Fig. 5E-G). Collectively, these data illustrate that Ch- exacerbates tau pathology in 3xTg-AD mice.

**Figure 5.**
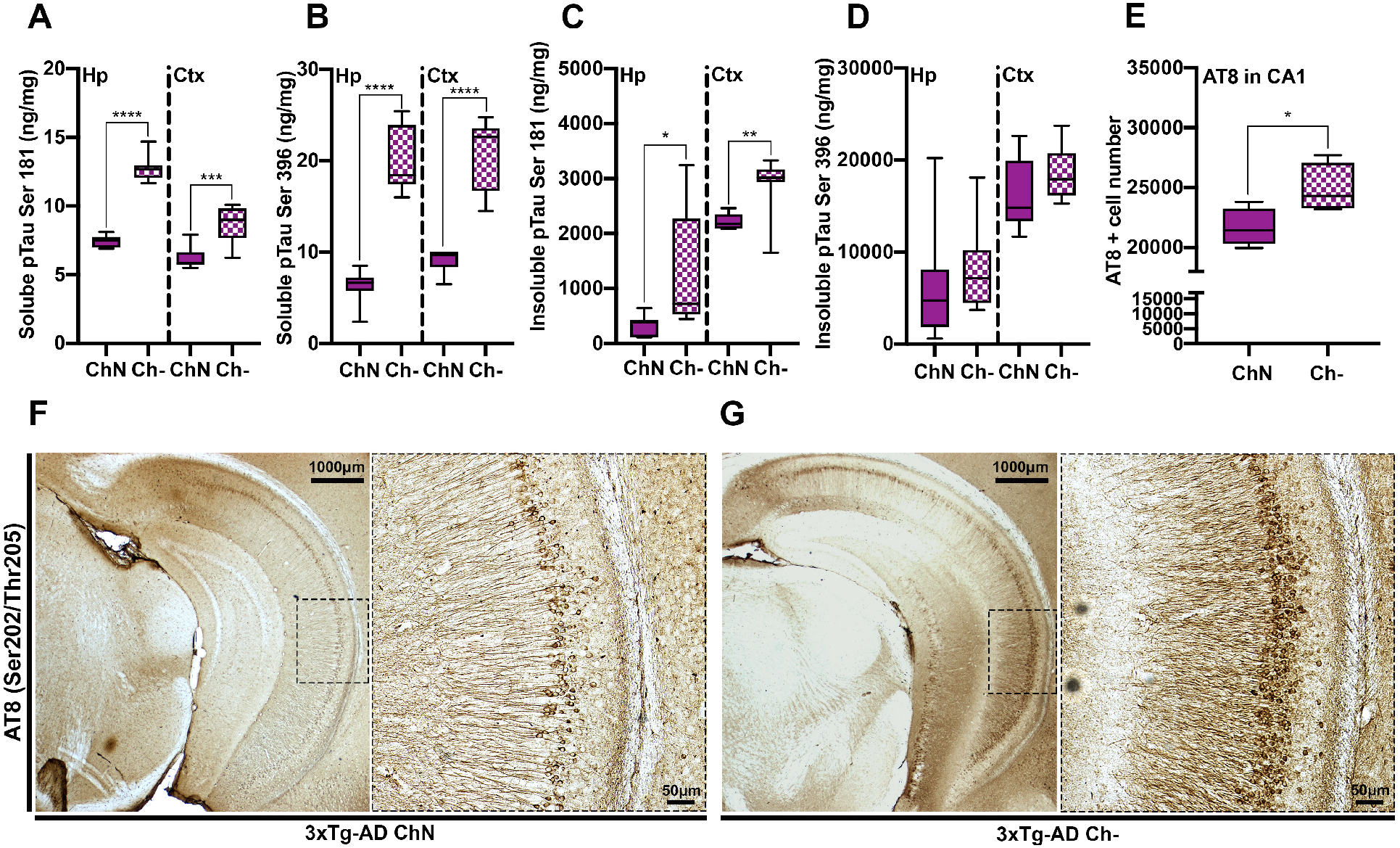
Ch- increases phosphorylation of pathological tau epitopes in the hippocampus and cortex of 3xTg-AD mice. **A**. Soluble levels of phosphorylated tau (pTau) at Serine (Ser) 181 were significantly elevated in the hippocampus (Hp; p < 0.0001) and cortex (Ctx; p = 0.0009) of 3xTg-AD Ch- mice. **B**. Soluble levels of pTau at Ser396 were significantly elevated in the Hp (p < 0.0001) and Ctx (p < 0.0001) of 3xTg-AD Ch- mice. **C, D**. Insoluble levels of pTau at Ser181 were significantly elevated in the Hp (p = 0.0442) and Ctx (p = 0.0052) of 3xTg-AD Ch- mice. No significant difference was observed for insoluble pTau at Ser 396 in the Hp and Ctx. **E**. AT8 (Ser202/Threonine (Thr)205)-positive cell numbers were significantly elevated in the CA1 region of the Hp of 3xTg-AD Ch- mice (p = 0.049). **F, G**. Photomicrographs of Ventral Hp sections stained for AT8 at 2.5x and 10x. For box plots, the center line represents the median value, the limits represent the 25^th^ and 75^th^ percentile, and the whiskers represent the minimum and maximum value of the distribution. * p < 0.05, ** p < 0.01, *** p < 0.001, **** p < 0.0001.

### Ch- alters protein networks associated with metabolic processing in NonTg hippocampi

To further understand how Ch- altered protein networks in the hippocampus, we performed LC-MS/MS coupled with LFQ and identified 3,544 proteins (Supplemental Fig. 2). We conducted two analyses comparing the hippocampal proteome of the NonTg ChN mice (n = 4/group) to that of the NonTg Ch- mice and the hippocampal proteome of the 3xTg-AD ChN mice to that of the 3xTg-AD Ch- mice (adj. p-val < 0.05, −1 > Log_2_ FC > 1; Supplemental Fig. 2). In the NonTg ChN vs. NonTg Ch- comparison, 86 differentially abundant proteins were identified, and in the 3xTg-AD ChN vs. 3xTg-AD Ch- comparison, 249 differentially abundant proteins were identified (Fig. 6A-C). Thirty-four proteins were commonly identified as differentially abundant due to Ch- in both the NonTg and 3xTg-AD hippocampi (Fig. 6C; Supplemental Fig. 2).

**Figure 6.**
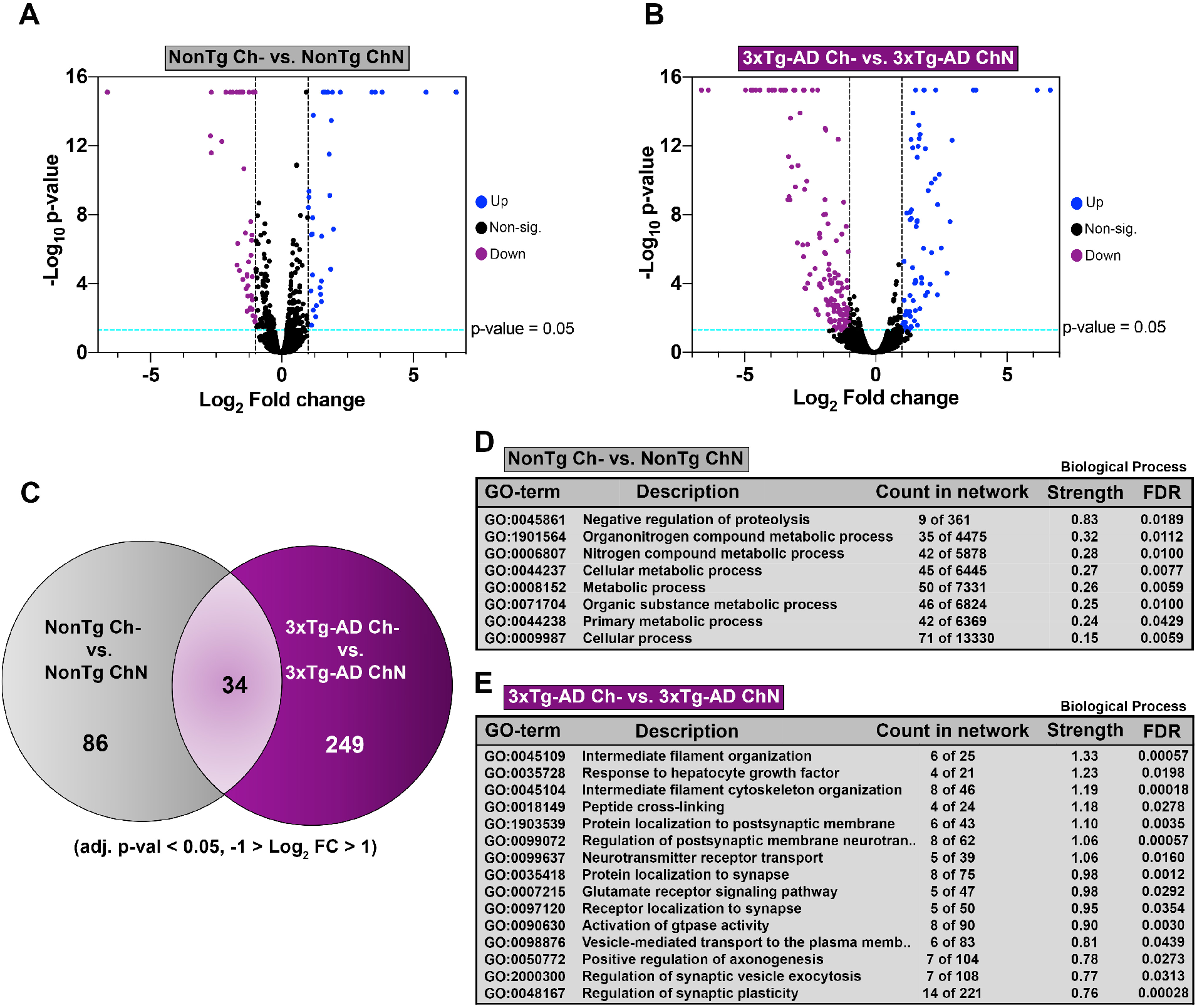
Ch- alters hippocampal protein networks in both NonTg and 3xTg-AD mice. **A, B**. Liquid chromatography tandem mass spectrometry followed by label free quantification (adj. p-val < 0.05, −1 > Log_2_ FC > 1) identified 86 differentially abundant proteins between NonTg Ch- and NonTg ChN hippocampi, and 249 differentially abundant proteins between 3xTg-AD Ch- and 3xTg-AD ChN hippocampi. **C**. Thirty-four proteins were commonly identified as differentially abundant due to Ch- in both NonTg and 3xTg-AD hippocampi. **D, E**. NonTg ChN vs. NonTg Ch- and 3xTg-AD ChN vs. 3xTg-AD Ch- gene ontology (GO) biological process classification analyses (for **E**, the top 15 biological processes are displayed based on strength of prediction).

Next, we performed gene ontology (GO) on the two sets of differentially abundant proteins to understand what altered biological processes and molecular functions were represented in these data sets. In the NonTg ChN vs. NonTg Ch- comparison, GO revealed changes in pathways associated with organonitrogen compound metabolic process, nitrogen compound metabolic process, and cellular metabolic process (Fig. 6D; full GO Supplemental Fig. 3). These findings corroborate choline’s well-documented role in metabolic processing (28).

### Ch- alters protein networks associated with microtubule function and postsynaptic membrane regulation in 3xTg-AD hippocampi

In the dataset of differentially abundant proteins between 3xTg-AD ChN and 3xTg-AD Ch- hippocampi, GO revealed changes in protein networks closely associated with AD pathology (Fig. 6E; full GO Supplemental Fig. 3). Altered biological processes included intermediate filament cytoskeleton organization, cytoskeleton organization, regulation of vesicle-mediated transport to plasma membrane, and vesicle-mediated transport in synapse. These pathways are particularly interesting because they are directly related to microtubule function, which is closely tied to tau pathology in AD (2). Hyperphosphorylation of tau results in its disassociation from the microtubules, disrupting microtubule stability and axonal trafficking (2). These data suggest that Ch- may also alter microtubule function in 3xTg-AD hippocampi and exacerbate microtubule dysfunction caused by tau pathogenesis. GO also revealed a series of altered biological processes associated with postsynaptic membrane regulation including protein localization to postsynaptic membrane, regulation of postsynaptic membrane neurotransmitter receptors, neurotransmitter receptor transport, protein localization to synapse, receptor localization to synapse, regulation of synaptic vesicle exocytosis, and regulation of synaptic plasticity. These findings corroborate previous work showing that adulthood dietary choline supplementation modulates the abundance of critical neuroreceptors in the 3xTg-AD hippocampus (8).

### Ch- alters protein networks associated with immune response and inflammation in NonTg plasma

To understand the mechanisms by which Ch- may contribute to systems-wide dysfunction, we next performed LC-MS/MS coupled with LFQ of the plasma proteome and identified 734 proteins (Supplemental Fig. 2). We performed two analyses, comparing the plasma proteome of the NonTg Ch- mice to that of the NonTg ChN mice (n = 4/group) and the plasma proteome of the 3xTg-AD Ch- mice to that of the 3xTg-AD ChN mice (adj. p-val < 0.05, −1 > Log_2_ FC > 1; Supplemental Fig. 2). In the NonTg Ch- vs. NonTg ChN comparison, 80 differentially abundant proteins were identified. In the 3xTg-AD ChN vs. 3xTg-AD Ch- comparison, 70 differentially abundant proteins were identified (Fig. 7A, B). Twenty-six proteins were commonly identified as differentially abundant due to Ch- in both the NonTg and 3xTg-AD plasma (Fig. 7C; Supplemental Fig. 2). GO of the list of differentially abundant plasma proteins in the NonTg ChN vs. NonTg Ch- comparison revealed that Ch- was implicated in dysregulation of biological processes related to inflammatory response, including acute-phase response, acute inflammatory response, response to inorganic substance, complement activation (alternative pathway), and defense response (Fig. 7D; full GO Supplemental Fig. 3). This is particularly interesting, as inflammation and immune dysfunction are relevant in a variety of diseases that affect all organs throughout the body, including AD (3–5, 29, 30).

**Figure 7.**
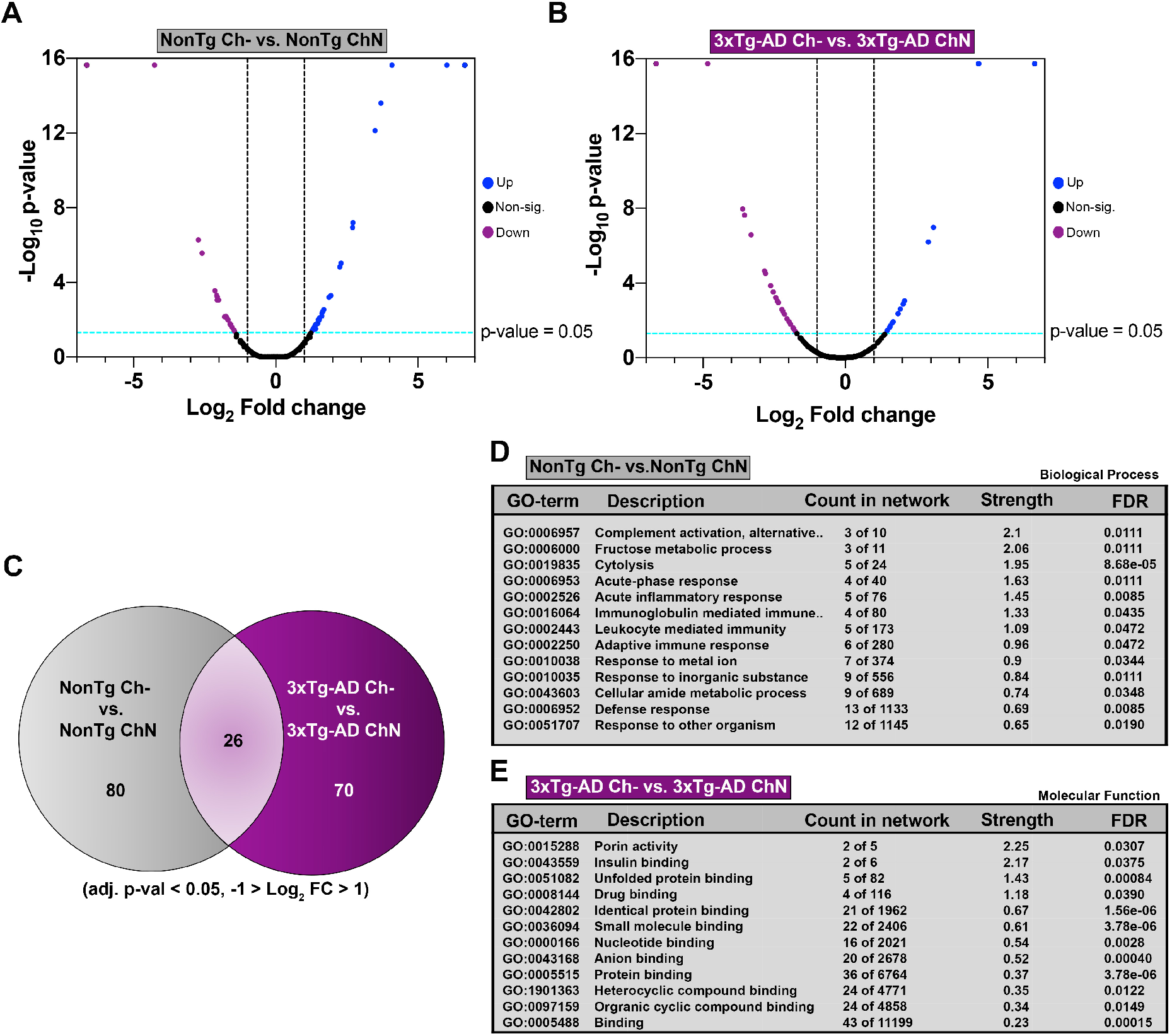
Ch- alters plasma protein networks in both NonTg and 3xTg-AD mice. **A, B**. Liquid chromatography tandem mass spectrometry followed by label free quantification (adj. p-val < 0.05, −1 > Log_2_ FC > 1) identified 80 differentially abundant proteins between NonTg Ch- and NonTg ChN plasma, and 70 differentially abundant proteins between 3xTg-AD ChN- and 3xTg-AD ChN plasma. **C**. Twenty-six proteins were commonly identified as differentially abundant due to Ch- in both NonTg and 3xTg-AD plasma. **D, E**. NonTg ChN vs. NonTg Ch- gene ontology (GO) biological process classification analysis and 3xTg-AD ChN vs. 3xTg-AD Ch- GO molecular function classification analysis.

### Ch- alters protein networks associated with insulin binding and porin activity in 3xTg-AD plasma

GO on the list of differentially abundant plasma proteins in the 3xTg-AD ChN and 3xTg-AD Ch- mice revealed that proteins associated with insulin binding were altered by Ch- (Fig. 7E). This is consistent with the glucose metabolism dysfunction we identified in the 3xTg-AD Ch- mice, suggesting that Ch- may modulate insulin binding proteins in the plasma, and thereby contribute to glucose metabolism impairments. GO also revealed that proteins associated with porin activity were altered by Ch-, including the voltage-dependent anion channel proteins VDAC-1 and VDAC-2 (Fig. 7E). These proteins and their roles in mitochondrial dysfunction have been previously associated with cell death and cognitive decline in AD patients (31).

## Discussion

Here, we showed that dietary choline deficiency (Ch-) throughout adulthood negatively impacts cellular and molecular function across a variety of systems. We found that Ch- led to motor impairments, weight gain, impaired glucose metabolism, cardiac pathology, and hepatic disease. Our measurements in plasma and brain tissue confirmed that Ch- resulted in significantly lower levels of choline. In the brain, we found that Ch- increased Aβ and pathological tau markers in both the hippocampus and cortex, two areas affected in AD (2). Our proteomic analysis of the hippocampus revealed that Ch- altered the expression of proteins networks related to AD pathogenesis including microtubule function and postsynaptic membrane regulation. In our plasma proteomic analysis, we found dysregulation of inflammatory-response and insulin-signaling related proteins, which corresponds to our physiological observations as well as well-known risk factors that link the peripheral system with the aging brain. Together, these data illustrate the importance of adequate dietary choline intake in promoting healthy aging across a variety of bodily systems and show that Ch- poses significant health risks for a large majority of the American population.

Choline is a precursor to ACh (9), the critical neurotransmitter in the neuromuscular junction that mediates voluntary motor function (32, 33). We observed motor deficits in the 3xTg-AD mice compared to NonTg mice, and in the Ch- mice compared to the ChN mice. We also show, for the first time, that 3xTg-AD ChN mice exhibit a reduction in plasma ACh compared to NonTg mice, and that Ch- resulted in further reduced plasma ACh, which may contribute to motor deficits. This is consistent with previous literature showing a direct relationship between dietary choline intake, ACh levels, and motor function (33). 3xTg-AD Ch- mice showed elevated Aβ pathology in the hippocampus and cortex; soluble and insoluble Aβ, Aβ oligomers, and Thioflavin S staining were all elevated in the 3xTg-AD Ch- mice compared to their ChN counterparts. Ch- also increased tau phosphorylation in the hippocampi and cortices of 3xTg-AD mice. Interestingly, this elevation of AD pathology did not correspond to worse performance in the MWM. Our previous work has shown that adulthood choline supplementation, in similar aged-mice, improved both AD pathology and cognitive performance in the MWM (8). Here, we hypothesize that further impairment of spatial learning and reference memory in the 3xTg-AD Ch- mice did not occur because of ceiling effects in MWM performance of the 3xTg-AD ChN mice (8). Our data support that adequate choline intake plays a critical role in protection against AD pathology.

Hyperinflammation, mitochondrial dysfunction, and glucose metabolism impairments are observed in AD (4, 6). These dysfunctions are not unique to the brain and are seen across the entire body. We observed that Ch- increased weight, impaired glucose metabolism and contributed to cardiac and liver pathology. Ch- mice had higher body weights than their ChN counterparts despite similar food consumption. Although 3xTg-AD mice typically show glucose metabolism impairments relative to NonTg mice (6), 3xTg-AD Ch- mice showed elevated glucose intolerance above that of their ChN counterparts. Further, the NonTg Ch- mice also showed glucose metabolism impairments, compared to their ChN counterparts, and in fact, NonTg Ch- mice had similar dysfunction to that seen in the 3xTg-AD ChN mice. Choline plays an important role in energy metabolism (15), suggesting that Ch- can contribute to the development of a diabetic state. Type 2 diabetes mellitus (T2DM) is a significant risk factor for AD (4, 6) and the two conditions are strongly interconnected (4). The 3xTg-AD Ch- mice not only showed the most severe weight gain and glucose metabolism impairments, but also had the highest pathological burden, suggesting that glucose metabolism impairments may have contributed to exacerbated AD pathology.

Ch- has been linked to clinical markers of cardiovascular disease (16), and there is an established link between cardiovascular disease and the development of dementias, including AD (5). Cardiovascular disease and AD have overlapping risk factors, including the *APOE4* gene, diabetes, a high fat diet, and a sedentary lifestyle (5). Moreover, cardiovascular issues increase the risk of AD through multiple channels including decreased cerebral blood flow (5). In the present study, we found that the 3xTg-AD phenotype and the Ch- diet led to cardiac hypertrophy, with the 3xTg-AD Ch- mice showing the most severe cardiac hypertrophy. We then measured the expression of genes associated with cardiac pathology (*Col1a1, Myh7*, and *Nppa)* and found that all groups except NonTg ChN had elevated mRNA expression of these markers. This illustrates that Ch- was sufficient to induce cardiac hypertrophy in NonTg mice, and further exacerbate cardiac hypertrophy in the 3xTg-AD mice, collectively highlighting that Ch- may increase risk for cardiac dysfunction.

Unsurprisingly, we detected markers of liver disease in the Ch- mice; NonTg mice showed NALFD as evidenced by hepatic steatosis, while 3xTg-AD mice had progressed to NASH, with evidence of inflammation and scarring alongside hepatic steatosis. Notably, while Ch- can induce NAFLD, work has shown that obesity, diabetes, and insulin resistance can also induce this phenotype, which was observed in both Ch- groups and may have also contributed to liver pathology (30). A healthy liver is critical for clearance of circulating Aβ in the periphery and hepatic disease reduces the clearance of peripherally-circulating Aβ (34). The peripheral clearance of Aβ by the liver helps reduce the accumulation of Aβ into plaques (34). We found that Aβ pathology was increased in the brains of 3xTg-AD Ch- mice, and failure to effectively clear Aβ due to liver disease may partially contribute to the increased burden, although future work is necessary to further interrogate this hypothesis.

Proteomics analysis of hippocampal tissue identified Ch- induced changes in key proteins linked to AD-related biological processes. The MAPT was altered in 3xTg-AD Ch- hippocampi, where 3xTg-AD Ch- mice showed reduced protein expression compared to the 3xTg-AD ChN mice. This is notable given the important role of healthy tau in microtubule stabilization, protein transport, synaptic plasticity and associated learning and memory (35) and highlights the effect of Ch- on this biological process. Proteomic analysis also revealed Ch- induced modulation of hippocampal protein networks associated with postsynaptic membrane regulation, suggesting that Ch- may also be linked to the synaptic dysfunction observed in AD. Interestingly neuroplastin (NPTN), a critical protein for long-term-potentiation at hippocampal excitatory synapses that is also linked to AD (36), was altered by Ch- in 3xTg-AD hippocampi. These proteins are important in learning and memory, which is often impacted in early AD (2). Moreover, proteins that modulate AMPA receptors (CACNG8, LRRTM1, FRRS1L), as well as NMDA receptors (GRIN2A, CLSTN1), were also altered by Ch- (37–41), suggesting that Ch- can contribute to postsynaptic membrane dysfunction in AD. Previous work has shown that choline, a precursor to ACh, modulates the expression of alpha7 nicotinic ACh receptor (α7nAChR) and Sigma-1R (σ1) R (8). Our results corroborate these previous findings that adulthood choline supplementation regulates the expression of postsynaptic receptors and suggest that choline alters postsynaptic receptor abundance through additional pathways outside of ACh. Altogether, these results provide significant evidence that Ch- exacerbates AD pathology and synaptic dysfunction.

Given the systems-wide dysfunction observed, we performed comparative proteomics on plasma from ChN and Ch- NonTg and 3xTg-AD mice. Interestingly, Ch- modulates key inflammatory and immune response pathways. More specifically, within the acute inflammatory response pathway, Ch- altered serum amyloid A 1 and 2 (SAA1 and SAA2, respectively) in both NonTg and 3xTg-AD mice. These two serum amyloid proteins are produced peripherally as an inflammatory response to environmental insult (42). SAA1 has been shown to prime microglia for ATP-dependent interleukin-1B release, which is associated with AD onset (42). These findings are particularly insightful, as they corroborate previous literature demonstrating that adulthood choline supplementation in an AD mouse model decreases disease-associated microglial activation (8). Previous literature also indicates that glial cell populations are responsive to SAA1 secretion in the blood and that SAA1 overexpression increases amyloid aggregation and glial activation in an AD mouse model (42, 43). This suggests that Ch- modulates SAA1/2 secretion in the plasma, causing glial hyperactivation, increased neuroinflammation, and even increased Aß aggregation. With respect to increased insulin resistance caused by Ch- in healthy aging and in AD, we find that a plasma protein network associated with insulin binding (containing insulin-degrading enzyme (IDE) and heat shock protein family D (HSPD1, also known as HSP60)) was altered due to Ch-. IDE degrades insulin, is directly related to insulin resistance, and also degrades Aß (4). IDE and its impact on insulin resistance have been linked to cognitive impairment (44). Additionally, there is growing evidence that HSPD1 modulates diabetes-induced inflammation and protects against Aß oligomer-induced synaptic toxicity (45, 46). This is particularly important given the large body of literature that demonstrates the negative effects of T2DM on brain insulin resistance, oxidative stress, and cognitive decline (4). Taken together, Ch- modulates key insulin binding proteins, which may contribute to the diabetes-like pathology observed in AD cases. Finally, mitochondrial dysfunction has emerged as another driver of AD pathology (4). Our results indicate that Ch- modulates two plasma proteins involved in porin binding, VDAC-1 and VDAC-2, that have also been linked to mitochondrial dysfunction and AD pathology (31, 47). This suggests that VDAC-1 and VDAC-2 modulation by Ch- may induce mitochondrial dysfunction in AD mouse models. VDAC-1 has been shown to mediate Aß toxicity in the brain, and its overexpression triggers cell death (31). Additionally, VDAC-1 interacts with several AD-relevant proteins, including pTau, Aß, and gamma-secretase (31). Interestingly, VDAC-1 serum expression has been strongly correlated with cognitive decline assessed in AD patients (48).

In conclusion, adequate choline intake is important for health across a variety of bodily systems, including the metabolic, cardiac, hepatic, and neurological systems. We found that Ch- led to pathological alterations across these systems in both NonTg and 3xTg-AD mice, suggesting that Ch- induces system-wide cellular and molecular dysfunction and increases the risk of AD across several pathogenic axes. While conducted in mice, if generalized to humans, these findings may help mitigate the estimated increase in prevalence of AD, illustrating the importance of adequate dietary choline intake throughout adulthood to offset disease occurrence for the general population.

## Materials and Methods

### Study design

This controlled laboratory experiment used 3xTg-AD and NonTg (C57BL6/129Svj) mice that were randomly assigned to one of two diets at three months of age; a standard laboratory chow diet (Envigo Teklab Diets, Madison WI) with normal choline levels (ChN; 2.0g/kg; #TD.180228), based on the human ADI or a laboratory chow choline deficient (Ch-; 0.0g/kg; #TD.110617; Supplemental Fig. 1A, B) diet. The start age for diet manipulation was selected due to choline’s important role in the developing brain (9). Sample sizes selected were equivalent to published work showing behavioral deficits and pathology in 3xTg-AD mice (6, 7). This resulted in four experimental groups (NonTg ChN, n = 20; 3xTg-AD ChN, n = 15; NonTg Ch-, n = 18; 3xTg-AD Ch-, n = 16). Behavioral assessment and endpoint tissue collection were performed at timepoints corresponding to known pathology in 3xTg-AD mice. Experimenters were blind to the groups. Exclusions were made when mice were unable to complete a behavioral test.

### Mice strain generation and housing

3xTg-AD mice were generated as previously described (6). 3xTg-AD mice harbor homozygous pathological mutations for the amyloid precursor protein (APP; Swedish), presenilin 1 (PSEN1; M146V), and microtubule associated protein Tau (MAPT; P301L) genes and were maintained in colonies by breeding homozygous 3xTg-AD mice with one another, on a C57BL6/129Svj hybrid background. C57BL6/129Svj mice were used as non-transgenic controls (NonTg). Female 3xTg-AD mice display consistent neuropathology while male 3xTg-AD mice do not, therefore consistent with published work, only female mice were used (6). All mice were kept on a 12-hour light/dark cycle at 23°C with *ad libitum* access to food and water. Mice were group housed, four to five mice per cage.

### Behavioral testing

At 10 months of age, mice underwent three days of rotarod testing (AccuScan Instruments Inc.) to assess motor ability as previously described (35). Mice were trained 6 trials/day for two days followed by a probe day. On training days, the rotation of the rod increased by 0.75rpm/s over 20s for a maximum speed of 15rpm. On the probe trials, the rod accelerated at a steady increase of 1 rpm/s for up to 90s. Mice were then tested in the Morris water maze (MWM) as previously described (7, 8). A circular pool with a platform 1cm under water obscured with white paint was used. All mice underwent 4 trials/day/5days for training. Location of the hidden platform remained constant, but the start location pseudo-randomly varied across trials. Mice were given 60s/trial to locate the hidden platform. Failure to reach the hidden platform resulted in guidance to its location. Twenty-four hours after the last training session, the platform was removed, and mice were returned to the MWM for 60s to assess spatial reference memory. A video camera recorded each trial. Data was analyzed via EthoVisionXT (Noldus Information Technology).

### Weight, food consumption and glucose tolerance (GTT) test

Weight was measured biweekly. A food consumption test was performed in 9-month-old mice as previously described (49). Mice were housed in 18 cages balanced for genotype and diet. Food was added at equal amounts on Day 1 and weighed every 24 hours for six days to assess intake, measuring food consumed per day / number of mice in a cage. A GTT was performed as previously described prior to euthanasia (6). Animals were fasted overnight for 16 hours, and baseline fasting glucose levels were taken. Animals then received 2 mg/kg glucose intraperitoneal (i.p.) injection and blood glucose was sampled from the tail using a TRUEtrack glucose meter and TRUEtrack test strips (Trividia Health) at 15, 30, 45, 60, 90, 120, and 150 minutes following the injection.

### Blood collection and plasma extraction

Mice were fasted for 16 hours, and blood was collected via the submandibular vein. 150 – 200 µL (≤ 10% of the subject’s body weight) of blood was collected and placed into EDTA lined tubes (BD K_2_EDTA #365974)) and inverted 8 times to assure anticoagulation. Tubes were kept on ice for 60-90 minutes and then centrifuged at 2200 RPM for 30 minutes at 4°C to separated phases. The top layer was collected and frozen at −80°C.

### Tissue harvesting and processing

Mice were euthanized at 12 months of age. One set of mice was perfused with 1X PBS, and had its brains and livers extracted and fixed in a glass vial filled with 4% paraformaldehyde for 48 hours. The remaining non-perfused mice had their brains extracted and their hippocampal and cortical tissue dissected, separately flash-frozen, and homogenized in a T-PER tissue protein extraction reagent supplemented with protease (Roche Applied Science) and phosphatase inhibitors (Millipore). Homogenized brain tissue was then centrifuged at 4°C for 30 mins, and the supernatant was decanted to be used as the soluble fraction for various assays. The pellet was homogenized in 70% formic acid solution and centrifuged at 4 ° C for 30 minutes to be used as the insoluble fractions for assays. Hearts were removed, weighed, and immediately flash frozen in dry ice. Heart total RNA was extracted from left ventricular extract using the RNeasy Mini Kit (Qiagen) as previously described (50). All qPCR probes were obtained from Integrated DNA Technologies.

### ELISA, western blots, choline, acetylcholine assays

We used commercially available ELISA kits (Invitrogen-ThermoFisher Scientific) to quantify soluble and insoluble levels of Aβ, and levels of phosphorylated Tau (pTau) at serine (Ser)181 and Ser396 as previously described (8). Choline and acetylcholine (ACh) levels were quantified using commercially available kits (Abcam, ab219944; Cell Biolabs, Inc., STA-603). Western blots were performed under reducing conditions as previously detailed (6) to probe for PEMT (dilution 1:1000, Thermo Fisher Scientific, PA5-42383) and loading control GAPDH (dilution 1:5000, Abcam, ab8245).

### Tissue sectioning and staining

Liver fixed tissue was sectioned at 20µm using a vibratome (Leica VT1000S), stained using a Hematoxylin and Eosin (H&E) kit (Abcam, ab245880), and imaged on a light microscope at 40X (Zeiss Axio Imager M2) to score for pathology as previously described (51). Brain hemispheres were sectioned into 50µm coronal sections using a vibratome and stored in specimen plates in PBS with 0.02% sodium azide. For Thioflavin S staining, tissue sections were incubated in filtered 1% aqueous Thioflavin-S for 10 minutes at room temperature, washed twice in 80% ethanol, once in 95% ethanol, and 3 times in ddH2O. Images were taken at 20x on a confocal microscope (Nikon A1R HD25). AT8 immunohistochemistry was performed as previously described (27) with the appropriate antibodies (1:1000 dilution, Thermo Fisher Scientific, catalog# MN1020).

### Unbiased stereology

Stereoinvestigator 17-software (Micro-BrightField, Cochester, VT) optical fractionator method was used to quantify AT8-positive CA1 cells in the hippocampus as previously described (27). Counts were performed at predetermined intervals; Grid size (*X* and *Y* = 158 μm), counting frame (*X* and *Y* = 50 μm), superimposed on the live image of the tissue sections. The sections were analyzed using a 63*x* × 1.4 PlanApo oil immersion objective. Gunderson score remained ≤ 0.08. Average tissue thickness was 26 μm. Dissector height was set at 22 μm, with a 2-μm top and 2-μm bottom guard zone. Seven sections were evaluated per animal. Bright field photomicrographs were taken with a Zeiss Axio Imager. The AT8 antibody penetrated the full depth of the section, allowing for the equal probability of counting all objects.

### Liquid chromatography tandem mass spectrometry (LC-MS/MS)

For LC-MS/MS, solubilized hippocampal mouse brain and plasma proteins were quantified (Thermo Fisher EZQ Protein Quantitation Kit or the Pierce BCA). Proteins were reduced with 50 mM dithiothreitol (Sigma-Aldrich) final concentration at 95°C for 10 minutes and alkylated with iodoacetamide (Pierce,40 mM final 30 minutes). Proteins were digested using 2.0 µg of MS grade porcine trypsin (Pierce) and peptides recovered using Protifi S-trap Micro Columns per manufacturer directions. Recovered peptides were dried via speed vac and resuspended in 30 µL of 0.1% formic acid.

All LC-MS analyses were performed at the ASU Mass Spectrometry Core Facility (cores.research.asu.edu/mass-spec/SOP). All data-dependent mass spectra were collected in positive mode using an Orbitrap Fusion Lumos mass spectrometer (Thermo Scientific) coupled with an UltiMate 3000 UHPLC (Thermo Scientific). One µL of peptide was fractionated using an Easy-Spray LC column (50 cm × 75 µm ID, PepMap C18, 2 µm particles, 100 Å pore size, Thermo Scientific) with an upstream 300µm x 5mm trap column. Electrospray potential was set to 1.6 kV and the ion transfer tube temperature to 300°C. The mass spectra were collected using the “Universal” method optimized for peptide analysis provided by Thermo Scientific. Full MS scans (375–1500 m/z range) were acquired in profile mode with the following settings: Orbitrap resolution 120,000 (at 200 m/z), cycle time 3 seconds and mass range “Normal”; RF lens at 30% and the AGC set to “Standard”; maximum ion accumulation set to “Auto”; monoisotopic peak determination (MIPS) at “peptide” and included charge states 2-7; dynamic exclusion at 60s, mass tolerance 10ppm, intensity threshold at 5.0e^3^; MS/MS spectra acquired in a centroid mode using quadrupole isolation at 1.6 (m/z); collision-induced fragmentation (CID) energy at 35%, activation time 10 milliseconds. Spectra were acquired over a 240-minute gradient, flow rate 0.250 uL/min as follows: 0-3 minutes at 2%, 3-75 minutes 2-15%, 75-180 minutes at 15-30%, 180-220 minutes at 30-35%, 220-225 minutes at 35-80% 225-230 at 80% and 230-240 at 80-5%.

### Label-free quantification (LFQ) and gene ontology (GO)

Raw spectra were loaded into Proteome Discover 2.4 (Thermo Scientific) and protein abundances determined using the Uniprot (www.uniprot.org) *Mus musculus* database (Mmus_UP000000589.fasta). Protein abundances were determined using raw files and were searched using the following parameters: Trypsin as enzyme, maximum missed cleavage site 3, min/max peptide length 6/144, precursor ion (MS1) mass tolerance at 20 ppm, fragment mass tolerance at 0.5 Da, and a minimum of 1 peptide identified. Carbamidomethyl (C) was specified as fixed modification, and dynamic modifications set to Acetyl and Met-loss at the N-terminus, and oxidation of Met. A concatenated target/decoy strategy and a false-discovery rate (FDR) set to 1.0% were calculated using Percolator (52). Accurate mass and retention time of detected ions (features) using Minora Feature Detector algorithm were then used to determine the area-under-the-curve (AUC) of the selected ion chromatograms of the aligned features across all runs and relative abundances calculated. GO analyses for biological process classification and molecular function classification were performed using STRINGv11.5 (Search Tool for the Retrieval of Interacting Genes/Proteins) as previously described (53).

### Statistical analyses

Two-way factorial Analysis of variance (ANOVA; for genotype and diet), were used to analyze the physiological and pathological experiments, followed by Bonferroni’s corrected post hoc tests when appropriate. Repeated measures ANOVA was used to analyze the behavior and GTT data. Student’s unpaired t-tests were utilized for comparison of 3xTg-AD mice. Levene’s test (to assess homogeneity of variance) revealed no significant effects necessitating the use of statistical tests other than the ones used. Significance was set at *p* < 0.05.

### Study approval

All animal procedures were approved in advance by the Institutional Animal Care and Use Committee of Arizona State University.

## Author Contributions

ND and JMJ: Determination of first co-authors was decided based on equal contribution to writing, data analysis and having performed experiments. AD: Animal studies, histology, edited the manuscript. WW: ELISA, statistical analysis, wrote the manuscript. PS: Liver pathology assessment, wrote the manuscript. OVE: Liver pathology assessment, wrote the manuscript. JS: Unbiased proteomics, edited the manuscript. AB: Animal studies, cardiac pathology, edited the manuscript. ST: Animal studies, edited the manuscript. IM: Data analysis, edited the manuscript. JKW: Data analysis, edited the manuscript. EAB: Cardiac pathology, edited the manuscript. CG: Experimental design, funding, cardiac pathology, edited the manuscript. TM: Experimental design, proteomic analysis, edited the manuscript. RV: Experimental design, funding, animal studies, wrote the manuscript. All authors read and approved the final manuscript.

## Acknowledgments

This work was supported by grants to R.V. from the NIH (R01 AG059627 and R01 AG062500).

## Data availability statement

Proteomic data have been deposited to the ProteomeXchange Consortium via the PRIDE partner repository.

## Supplementary Figure legends

**Supplemental Figure 1. A-B**. Datasheets of the choline normal (ChN) and deficient (Ch-) diets.

**Supplemental Figure 2**. Total proteins identified from hippocampal and plasma samples, and comparisons between NonTg Ch- vs. NonTg ChN and 3xTg-AD ChN vs. 3xTg-AD Ch-.

**Supplemental Figure 3**. Complete outputs from gene ontology (GO) analyses for biological process classification and molecular function classification of differentially abundant proteins in hippocampal (Hp) and plasma (pl.) samples.

