## Supplementary figures and images for "Dietary choline intake is necessary to prevent systems-wide organ pathology and reduce Alzheimer’s disease hallmarks"

### Supplemental Figure 1

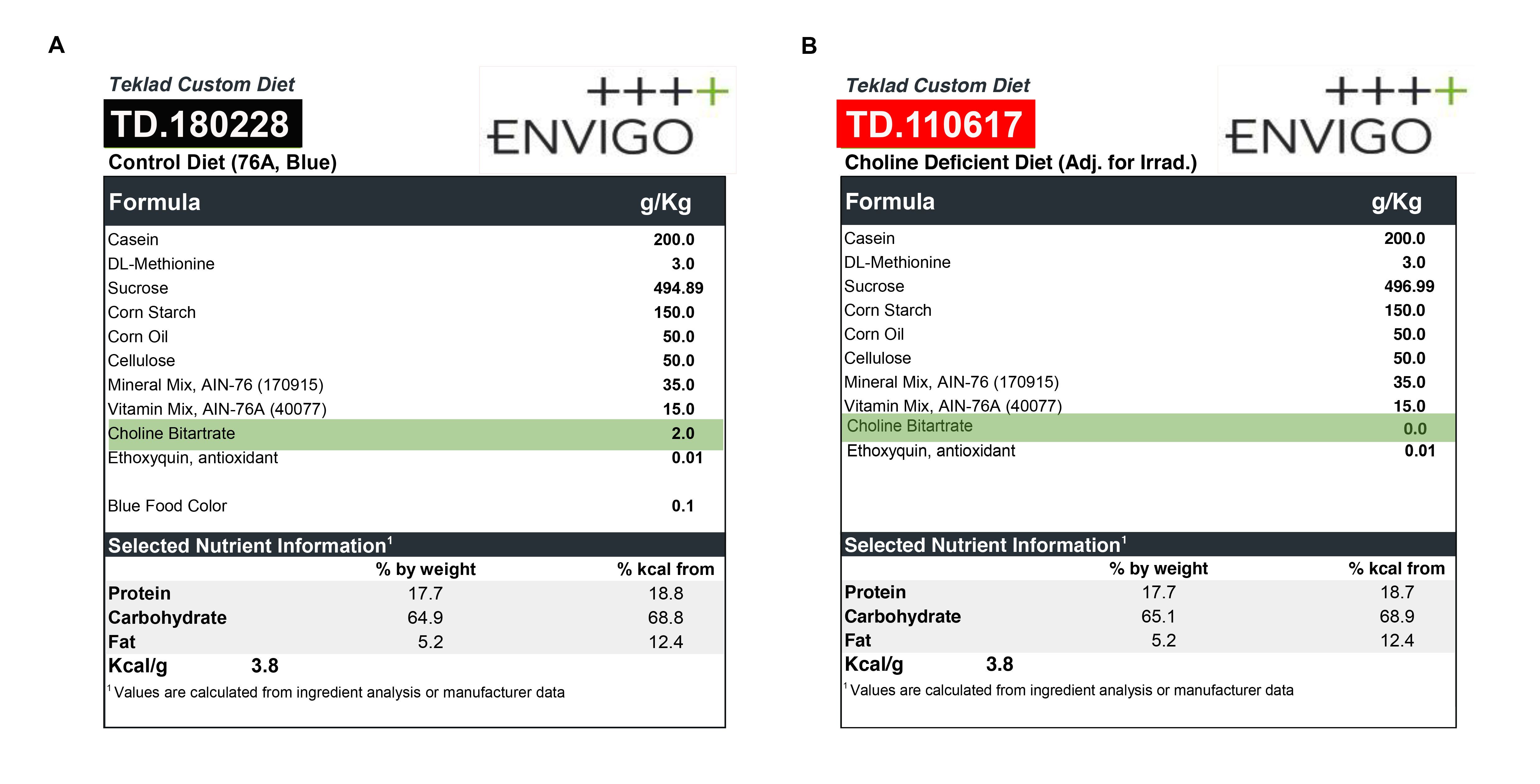
